# Diving Deep into Fish Bornaviruses: Uncovering Hidden Diversity and Transcriptional Strategies through Comprehensive Data Mining

**DOI:** 10.1101/2023.08.23.554433

**Authors:** Mirette Eshak, Dennis Rubbenstroth, Martin Beer, Florian Pfaff

## Abstract

Recently, we discovered two novel orthobornaviruses in colubrid and viperid snakes using an *in silico* data mining approach. Here, we present the results of a screening of more than 100,000 nucleic acid sequence datasets of fish samples from the Sequence Read Archive (SRA) for potential bornaviral sequences. We discovered the potentially complete genomes of seven bornaviruses in datasets from osteichthyans and chondrichthyans. Four of these are likely to represent novel species within the genus *Cultervirus*, and we propose that one genome represents a novel genus within the family of *Bornaviridae*. Specifically, we identified sequences of Wǔhàn sharpbelly bornavirus (WhSBV) in sequence data from the widely used grass carp liver and kidney cell lines L8824 and CIK, respectively. A complete genome of Murray-Darling carp bornavirus (MDCBV) was identified in sequence data from a goldfish (*Carassius auratus*). The newly discovered little skate bornavirus (LSBV), identified in the little skate (*Leucoraja erinacea*) dataset, contained a novel and unusual genomic architecture (N-Vp1-Vp2-X-P-G-M-L), as compared to other bornaviruses. Its genome is thought to encode two additional open reading frames (tentatively named Vp1 and Vp2), which appear to represent ancient duplications of the gene encoding for the viral glycoprotein (G). The datasets also provided insights into the possible transcriptional gradients of these bornaviruses and revealed previously unknown splicing mechanisms.

## INTRODUCTION

The family *Bornaviridae* belongs to the order *Mononegavirales* and includes viruses that are considered zoonotic and can cause severe disease in humans, such as Borna disease virus 1 (BoDV-1) [1] and the variegated squirrel bornavirus 1 (VSBV-1) [2]. Other members are of veterinary interest because they can cause severe disorders in birds, such as parrots [3]. Taxonomically, the family *Bornaviridae* currently consists of the three genera *Orthobornavirus, Carbovirus* and *Cultervirus* [4]. Of these, the orthobornaviruses have the widest so far known host spectrum and have been identified in birds, reptiles and mammals [5]. Carbo- and culterviruses have up to now only been identified in reptiles and fish, respectively [5–8]. The genus *Cultervirus* currently comprises a single virus that has been discovered in fish (Wǔhàn sharpbelly bornavirus [WhSBV], species *Cultervirus hemicultri*) [5, 8]. Partial genome sequences of another cultervirus, Murray-Darling carp bornavirus (MDCBV), have recently been published, but its classification is still pending [7].

The genome of bornaviruses consists of an approximately 9 kb non-segmented and single-stranded RNA molecule of negative polarity (−ssRNA) [9]. Typically, six viral proteins are encoded by the viral genome: nucleoprotein (N), accessory protein X, phosphoprotein (P), matrix protein (M), glycoprotein (G) and the large protein (L) containing an RNA-directed RNA polymerase domain [5, 9]. The open reading frames (ORF) encoding these viral proteins are arranged in two known genomic architectures: i) 3’-N-X-P-<u>M-G</u>-L-5’ (genus *Orthobornavirus*) and ii) 3’-N-X-P-<u>G-M</u>-L-5’ (genera *Carbovirus* and *Cultervirus*). Bornaviral replication and transcription occur in the nucleus of infected cells [10] and multiple viral transcripts are produced using conserved transcription initiation and termination sites [11]. Atypically for mononegaviruses, bornaviruses use alternative splicing in order to control and diversify their transcriptional capacity [12, 13].

Recently, we used an *in silico* data mining approach based on ‘*Serratus*’ [14] in order to screen for traces of potential bornaviruses hidden in archived sequence data from public nucleic acid sequences databases, such as the Sequence Read Archive (SRA) [15]. The SRA stores raw nucleic acid sequence reads from next-generation sequencing runs from multidisciplinary research experiments, along with extensive metadata. In these archived sequencing reads, we identified and characterised two potential novel orthobornaviruses of colubrid and viperid snakes: Caribbean watersnake bornavirus (CWBV) and Mexican black-tailed rattlesnake bornavirus (MRBV), in datasets from a Caribbean watersnake (*Tretanorhinus variabilis*) and a Mexican black-tailed rattlesnake (*Crotalus molossus nigrescens*), respectively [15].

In the present study, we extended the search for previously undetected bornaviruses by screening 116,082 transcriptomic datasets from fish samples from the orders Osteichthyes and Chondrichthyes and identified seven bornavirus genomes.

## MATERIAL AND METHODS

### Selection of datasets

We generated a list of datasets using the European Nucleotide Archive (ENA) Browser advanced search portal (https://www.ebi.ac.uk/ena/browser/advanced-search) and selected the data type ‘raw reads’ using the search query:

~~~
(tax_tree(1476529) OR tax_tree(7777) OR tax_tree(7898) OR tax_tree(7878)) AND
(library_source=“METATRANSCRIPTOMIC” OR library_source=“TRANSCRIPTOMIC SINGLE CELL”
OR library_source=“VIRAL RNA” OR library_source=“TRANSCRIPTOMIC”)
~~~

Specifically, this search included the taxonomic units of jawless vertebrates (Cyclostomata; NCBI:txid1476529), cartilaginous fishes (Chondrichthyes; NCBI:txid7777), ray-finned fishes (Actinopterygii; NCBI:txid7898) and lungfish (Dipnomorpha; NCBI:txid7878). We further restricted the search to RNA-derived datasets from (meta-) transcriptomic or viral RNA sequencing experiments.

### Data mining of raw reads

In order to identify even single reads within the selected datasets that may be related to bornaviruses, we developed the bioinformatics pipeline ‘SRAminer’. The ‘SRAminer’ pipeline is based on *snakemake* [16], is multi-threading and can be run in any Linux-like environment. The code for ‘SRAminer’ and detailed instructions on how to use it can be found at: https://gitlab.com/FPfaff/sraminer.

A simplified workflow of the pipeline includes the steps (i) download, (ii) blast and (iii) report: (i) A subset of reads from each dataset is downloaded using *fastq-dump* (v3.0.3; SRA Toolkit). Typically, a subset of 100,000 to 1,000,000 reads is sufficient to identify datasets containing sequence reads of interest. (ii) Using *diamond blastx* (v2.0.15; [17]), the subset of reads is then searched against a user-provided protein database. In this case, we selected and obtained the protein sequences from all available members of the family *Bornaviridae* from NCBI. (iii) If at least a single read matches the search criteria, additional metadata for this dataset is obtained using *ffq* (0.0.4; [18]) and the results are summarised into individual reports using *R* [19] and *R markdown* [20].

### Further raw read processing

After an initial screening of subsets of each 100,000 reads using SRAminer, the most promising datasets were selected based on the number of reads matching the blastx search criteria and the inferred theoretical number of reads in the full dataset. We then downloaded the full datasets of these most promising SRA entries using *parallel-fastq-dump* (v0.6.7; [20]) and trimmed them for low quality regions and adapter contamination using *TrimGalore!* (v0.6.10; [21]) running in automatic mode. The trimmed reads were then used for *de novo* assembly with *SPAdes* genome assembler (v3.15.5; [22]) running in --rna mode. The resulting transcripts/contigs were then searched against the representative bornavirus protein database using *diamond blastx* (v2.0.15; [17]). Transcripts/contigs matching the search criteria were selected and imported into Geneious Prime (v2021.0.1) for further characterisation. Final genomes were additionally screened for any vector or adapter contamination using the NCBI VecScreen suite (https://www.ncbi.nlm.nih.gov/tools/vecscreen). To verify the nature of the sampled organism, we selected all contigs from the assembly that matched the mitochondrial cytochrome B gene (*MT-CYB*) and submitted them to the NCBI blastn suite (https://blast.ncbi.nlm.nih.gov/Blast.cgi).

### Genomic characterization

Potential ORFs were predicted using the Geneious Prime (v2021.0.1) ‘Find ORFs’ function and for identification the deduced amino acid sequences were searched against the non-redundant blast database (nr) using the NCBI blastp suite. The trimmed raw reads were mapped back to the respective potential viral genome using the Geneious Prime (v2021.0.1) generic mapper (options: medium sensitivity; find structural variants, short insertions, and deletions of any size) in order to visualise transcriptional profiles and potential splice junctions. Potential transcription initiation and termination sites were predicted based on sequence similarity to known bornaviral signal sequences [11]. They were further verified by manual inspection of the read coverage at these positions (e.g. transition to poly(A) at the termination sites). In addition to visual inspection of the potential transcription start and termination sites, we used MEME (v5.5.2) to discover conserved motifs.

### Genomic classification

For the phylogenetic characterisation of potential bornavirus genomes, we used amino acid alignments based on the predicted and translated N, G and L genes. The amino acid sequences of these genes were individually aligned with 19 reference sequences using Muscle (v3.8.425). The reference viruses were selected in order to represent all ICTV-accepted species of the family *Bornaviridae* (n=12) as well as viruses below the species level (n=7). The individual alignments were then concatenated into a single alignment and IQ-TREE (v2.2.2.6) was used to infer the phylogenetic relationships. Specifically, a partitioning model (-Q) was used that allowed for individual substitution models and evolutionary rates in each partition. The substitution model was selected automatically (-m MFP+MERGE) and branch support was assessed using the ultrafast bootstrap (-bb) and SH-aLRT tests (-alrt) with each 1,000,000 replicates each.

In addition, the Pairwise Sequence Comparison (PASC) [23, 24] was used to classify the potential bornaviral genomes within the family *Bornaviridae*. PASC is based on pairwise global nucleotide sequence alignments along the entire viral genome using a Blast-based approach.

For the prediction of the potential transmembrane domains (TM) along with the signal peptide (SP) and cleavage sites (CS) within the G protein of the new genomes, we used DeepTMHMM (pybiolib, version 1.1.944 [25]) and ProP-1.0 [26], respectively.

## RESULTS

### Data mining

During data mining, we analysed subsets of 116,078 raw transcriptomic SRA datasets from fish (jawless vertebrates, cartilaginous fish, ray-finned fish and lungfish; see Supplementary Table S1). In 72 of the 116,078 SRA datasets, we found at least one single read that matched one of the bornavirus protein references. For all of these 72 datasets, a *de novo* assembly of all available data was performed and the resulting contigs were scored (see Supplementary Table S2). In 8 of the *de novo* assembled datasets, we identified endogenous bornavirus-like elements (EBLs), which were not further analysed. In 4 datasets, we identified complete genomes from members of the viral family *Chuviridae*. In a further 15 datasets, none of the resulting contigs showed any sequence similarity to the bornavirus reference database. In 44 *de novo* assembled SRA datasets, full or nearly full-length bornaviral genomes were identified. As some of these SRA datasets represented either different organ samples from the same animal or multiple replicates belonging to a single study or were based on the very same cell line, we selected only representative genomes for further characterisation.

As a result, 7 complete and unique bornaviral genomes were assembled from SRA datasets SRR10323915, SRR6207428, SRR1299086, SRR13236436, SRR9592747, SRR17661348, and SRR17441645 (**Table 1**). The *MT-CYB* sequences assembled from each of these datasets matched those of the specified sampled organisms (Supplementary Table S3).

**Table 1:**
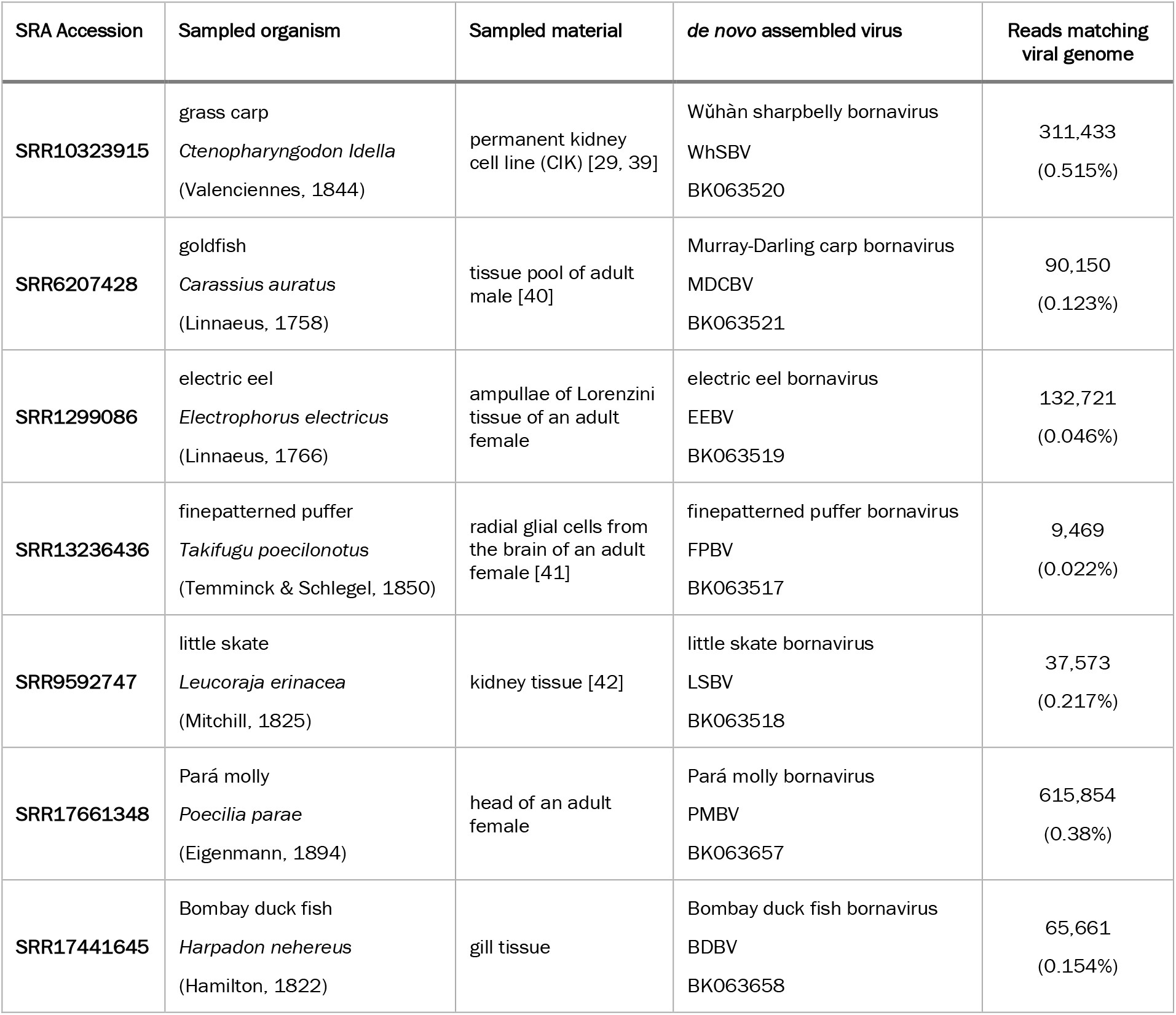
Summary of SRA datasets that were selected for *de novo* assembly of complete bornaviral genomes.

### Taxonomic relationship and classification

Phylogenetic analysis of the predicted viral proteins N, G, and L revealed that the potential bornaviruses clustered with viruses of the genus *Cultervirus*, represented by WhSBV (NC_055169) and MDCBV (MW645025-7), rather than carbo-or orthobornaviruses (Figure 1).

**Figure 1:**
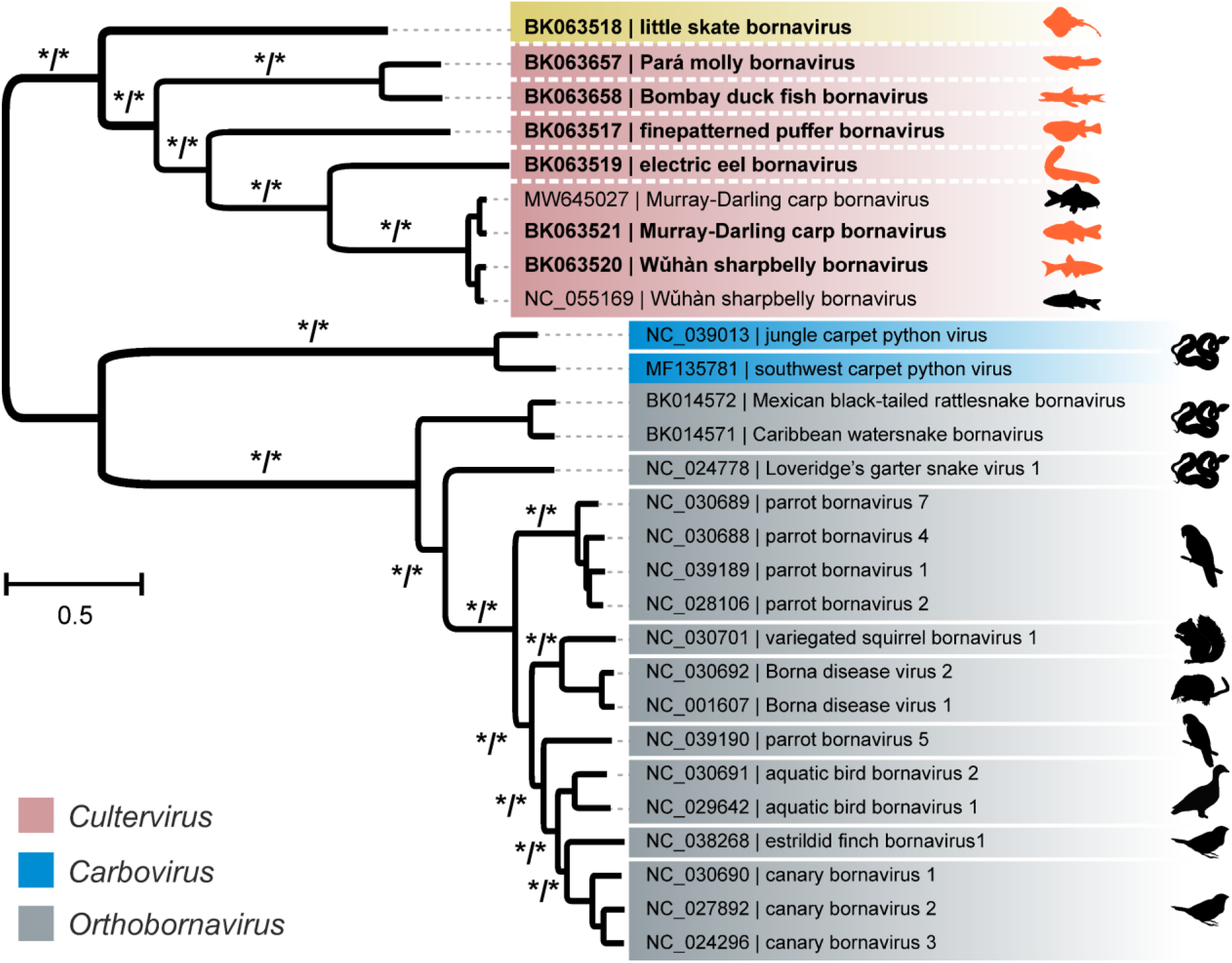
Phylogenetic relationships within the family *Bornaviridae*. The maximum-likelihood tree was based on the concatenated amino acid sequence alignments of the viral proteins N, G and L of the newly identified potential fish bornaviruses (bold) together with representative members of the genera *Cultervirus* (red), *Carbovirus* and *Orthobornavirus*. White lines indicate separate virus species. The silhouettes represent typical host organisms of previously published bornaviruses or the reported sampling source (highlighted) of the viral genomes identified in this study. The tree was constructed using IQ-TREE (version 2.2.2.3), an optimal partition model and statistical support with 1 million replicates each for ultrafast bootstrap and SH-aLRT test. Statistical support is shown for main branches using the format [ultrafast bootstrap/SH-aLRT]. Asterisks indicate statistical support ≥ 90% and ≥ 90% for ultrafast bootstrap and SH-aLRT, respectively.

Specifically, the full genome derived from a dataset derived from the grass carp kidney cell line CIK (SRR10323915) had 87.9% nucleotide identity to WhSBV (NC_055169) and was therefore considered to be a variant of WhSBV. We identified the nearly identical WhSBV genome sequence in 36 SRA datasets, all derived from RNA sequencing of either grass carp kidney (CIK; n=10) or liver (L8824; n=26) cell lines (Supplementary Table S4).

In contrast, the full bornavirus genome from a goldfish tissue pool dataset (SRR6207428) showed 99.5% nucleotide identity to partial sequences of MDCBV (MW645025-7). This sequence can therefore be considered the be the first complete genome of MDCBV.

Additional bornaviral sequences from Bombay duck fish (SRR17441645), electric eel (SRR1299086), Pará molly (SRR17661348), finepatterned puffer (SRR13236436), and little skate (SRR9592747) formed distinct taxonomic units. Hence, we tentatively named these potential viruses based on the origin of the underlying sampling material: Bombay duck fish bornavirus (BDBV; BK063658), electric eel bornavirus (EEBV; BK063519), Pará molly bornavirus (PMBV; BK063657), finepatterned puffer bornavirus (FPBV; BK063517), and little skate bornavirus (LSBV; BK063518). BDBV, EEBV, PMBV, and FPBV maintained between 42% and 66% PASC identity to the known culterviruses WhSBV and MDCBV and to each other (Supplementary Figure S1). At 65.8%, BDBV and PMBV were more closely related to each other than to any other virus. LSBV showed the greatest genetic divergence, with PASC identities ranging from 38.2% to 39.9% relative to all other viruses.

### Genome architecture

The genome architecture of MDCBV, BDBV, EEBV, PMBV, and FPBV was analogous to that of known culter- and carboviruses, characterised by the arrangement of genes as 3’-N-X/P-G-M-L-5’ (**Figure 2**). The identified grass carp WhSBV variant, as well as the goldfish MDCBV variant, closely resembled the WhSBV reference NC_055169 in structure and length (8,989 – 8,990 nt). In contrast, BDBV, EEBV, PMBV, and FPBV had genome lengths of 9,110, 9,148, 9,324, and 9,397 nt, respectively. Notably, the genome structure of LSBV differed from the other bornaviral genomes, as it was significantly longer, spanning 11,090 nt, and contained two additional ORFs designated viral proteins 1 and 2 (Vp1 and Vp2): 3’-N-Vp1-Vp2-X/P-G-M-L-5’.

**Figure 2:**
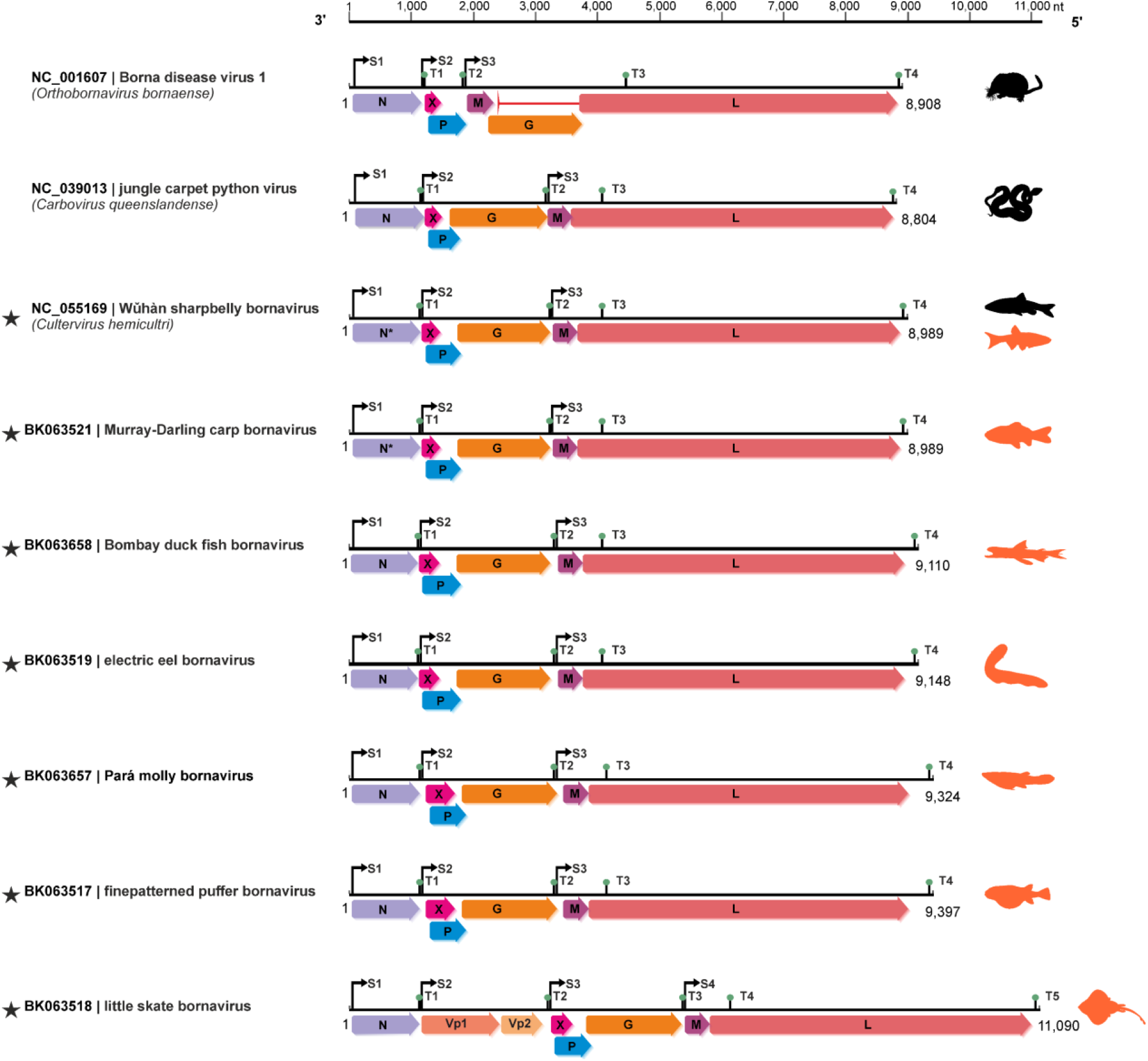
Genome architectures of current and potential novel bornaviruses. Representative overall genome organisations are shown for representative bornaviruses along with the potential novel viruses (star). The (predicted) open reading frames (ORF) are shown as arrows together with the predicted transcription start (S) and transcription termination (T) sites. For each of the genomes, the potential hosts/sources for each virus are shown. Note the different genomic arrangements: 3’-N-X-P-M-G-L-5’ (genus *Orthobornavirus*) and 3’-N-X-P-G-M-L-5’ (genera *Carbovirus* and *Cultervirus*). The little skate bornavirus shares the genomic structure of carbo- and culterviruses, but encodes two additional predicted ORFs: 3’-N-Vp1-Vp2-X-P-M-G-L-5’.

### Transcriptional profiles, motifs and alternative splicing

The transcriptional profiles and splice sites of the discovered bornavirus genomes were investigated by aligning/mapping the corresponding raw sequence data to the *de novo* assembled genomes (**Figure 2**). The observed sequence coverage was not uniform across the genomes and abrupt increases or decreases were observed within some of the potential intergenic regions. These changes in genome coverage colocalised with predicted transcription start and termination motifs. Specifically, the predicted start sites were characterised by a large increase in read coverage, whereas the termination sites correlated with decrease in read coverage and the presence of reads transitioning to poly(A) at the respective termination site. The respective positions of these predicted regulatory sites were highly conserved between the different viruses. In detail, start sites were present immediatly upstream of the N, X/P, and M ORFs. The potential termination sites were located downstream of the N, G and L ORFs. An additional termination site T3 was present within the L ORF (**Figure 3)**.

**Figure 3:**
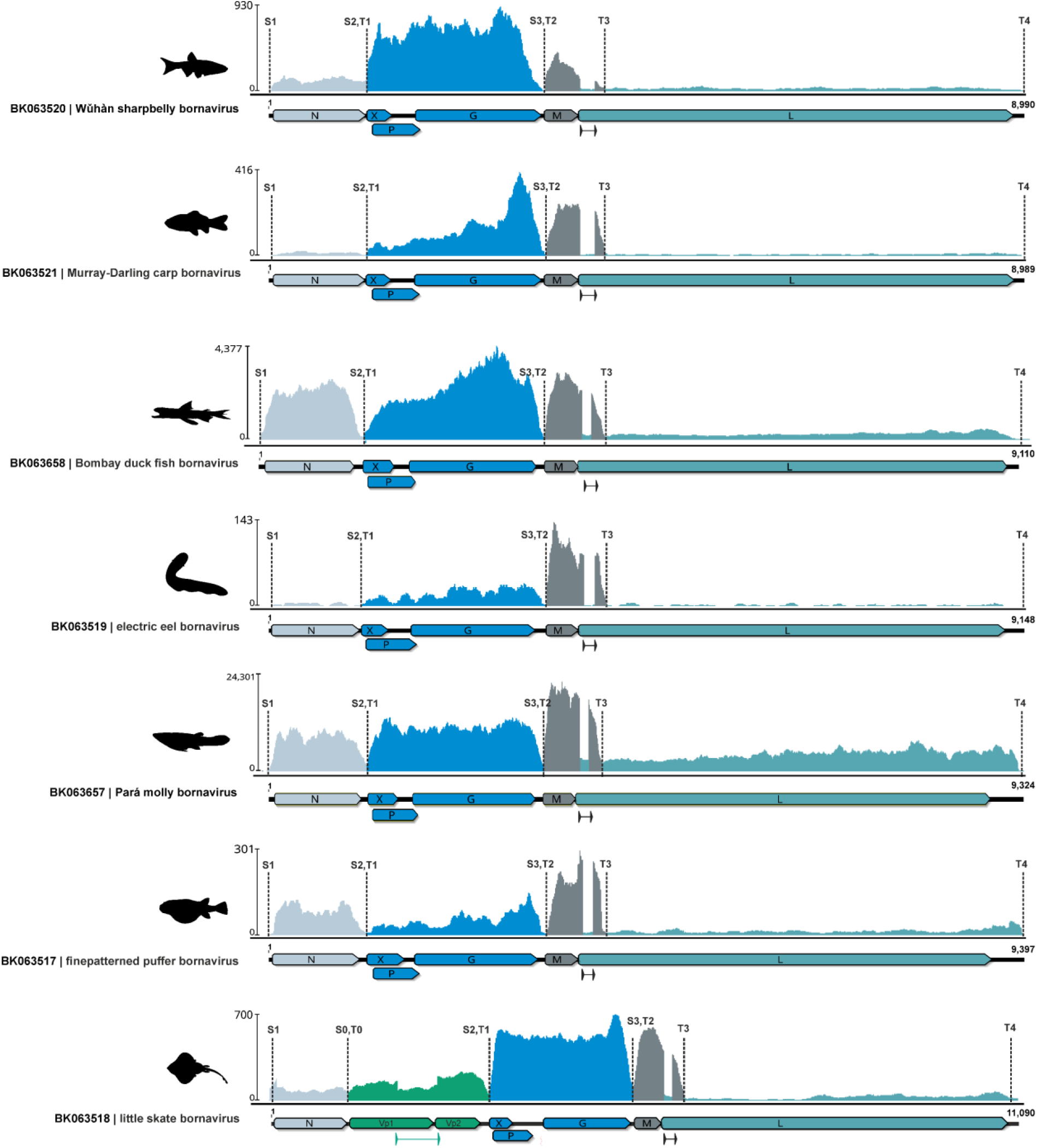
Transcriptional profiles of novel bornaviruses. Raw reads were mapped to the novel bornavirus genomes and the coverage was plotted. Open reading frames (ORFs) are shown as arrows and predicted transcription start (S) and transcription termination (T) motifs are indicted as dashed lines. S and T sites collocate with large increases and decreases in coverage, respectively. Regions that with similar coverage and are bordered by S and T sites were considered to represent individual RNA transcripts. These viral transcripts and their corresponding ORFs are highlighted in different colours. Alternative splicing was detected within in all viruses for potential M transcript (intron shown as line arrow). In addition, a potential intron was identified in the bicistronic transcript encoding Vp1 and Vp2 of little skate bornavirus.

Genomic regions that showed homogeneous coverage and were flanked by adjacent start and termination sites were interpreted as belonging to the same viral RNA transcripts or mRNA (**Figure 3**). The overall pattern of viral transcription was highly conserved among all fish bornaviruses analysed. In detail, the N protein appeared to be expressed from a monocistronic mRNA, whereas X/P and G were expressed from a polycistronic mRNA. The M and L transcripts appeared to share a single transcription start site (S3), but their expression levels were very different, with L being expressed at low levels and M at relatively high levels.

Interestingly, LSBV showed an additional start and termination site, that were located adjacent to the hypothetical ORFs of Vp1 and Vp2, suggesting that both proteins may be expressed from a bicistronic mRNA. An additional intron was identified between the Vp1 and Vp2 ORFs at nucleotide positions 1,868-2,476, which would result in an in-frame hybrid of the Vp1 and Vp2 ORFs, tentatively named Vp3 (see results below).

In addition, we identified an alternative splice site at the beginning of the L ORF, that was present in all viruses analysed. The ideentified splice site was supported by multiple reads missing the intronic sequence. The intron had a size of 110-176 nt and was located 23-53 nt downstream of the M ORF stop codon. The coverage depth of the unspliced RNA was comparable to that of the L ORF, while the spliced RNA had a coverage comparable to that of the M ORF (**Figure 3**). It could be speculated that M is expressed from an RNA that undergoes alternative splicing and uses the T3 transcription termination site located within the L ORF (**Figure 4A)**. The viral RNA for L on the other hand is expressed from the same S3 transcription start as M but does not undergo splicing and uses a the T4 termination site. The intronic sequences of all viruses analysed showed the canonical dinucleotides GU and AG for donor and acceptor sites, respectively (**Figure 4A**).

**Figure 4:**
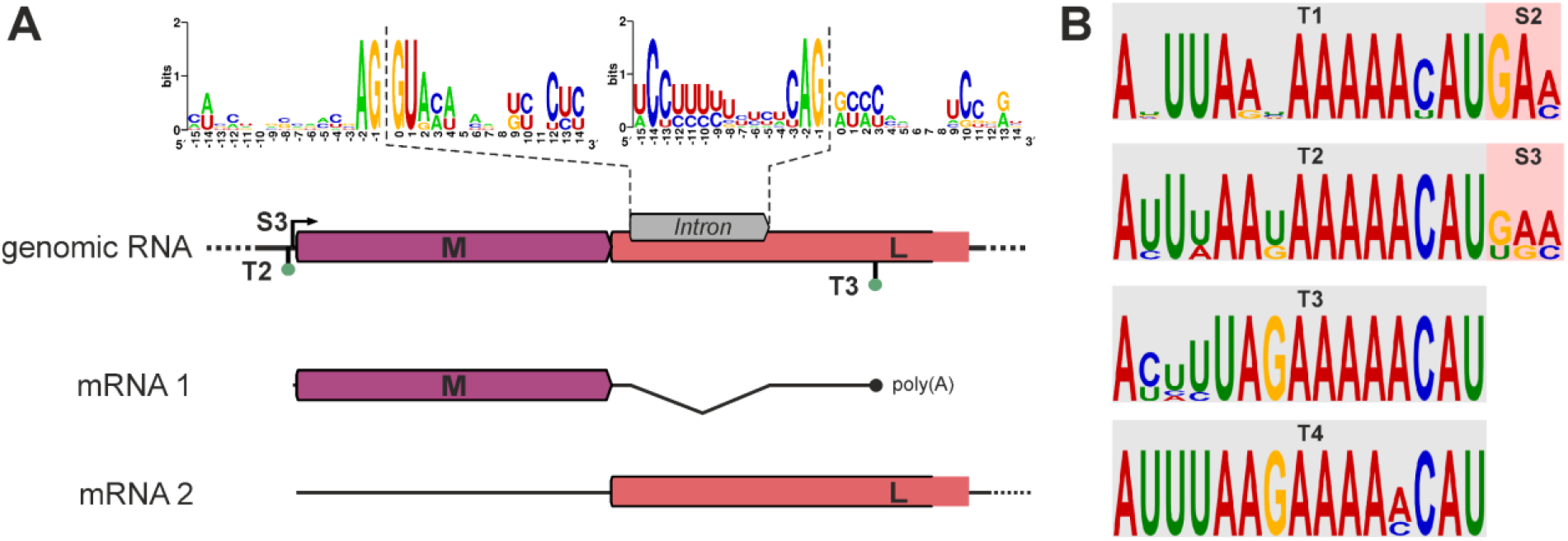
Conserved splicing mechanisms and transcriptional motifs among fish bornaviruses. (A) Alternative splicing was detected in the M/L ORF region for all viruses analysed. The genomic arrangement in this region is shown with ORFs indicated by arrows. The potential intron is shown as an arrow within the L ORF region. The sequence motifs of the splice acceptor and donor sites from all analysed viruses are shown and the dashed lines indicate the position of the splicing. The canonical GU/AG splice site is present in all viruses analysed. Two possible mRNAs are shown: The spliced mRNA contains only the M ORF and is terminated at the T3 site, while the unspliced mRNA will contains the full L ORF. (B) The conserved motifs of the transcription termination and start sites of the analysed viruses are shown. Note that T1/S2 and T2/S3 are directly adjacent to each other.

Motif prediction revealed conserved sequence patterns for transcription termination and start sites (**Figure 4B**). The termination sites T1-4 shared the conserved nucleotide sequence pattern `‘AYUUWAKAAAAACAU’`, whereas the start sites S1, S2 and S3 shared the conserved nucleotide sequence pattern `‘GAM’`. S2 and S3 were immediately adjacent to T1 and T2, respectively.

### LSBV Vp1 and Vp2 are homologues of the glycoprotein G

When analysed by pairwise alignment, the hypothetical viral proteins Vp1 and Vp2 of LSBV shared amino acid similarity with the glycoprotein G of LSBV (**Figure 5A**). In detail, the pairwise amino acid identity between the G and Vp1 was 28%, between the G and Vp2 it was 21% and between Vp1 and Vp2 it was 41% (**Figure 5B**). While Vp1 shares the N-terminus of the G protein, it lacks the C terminus. Vp2 shares only the central region of the G protein and lacks both, the respective N- and C-terminal regions of G. Both, Vp1 and Vp2, have no detectable transmembrane domain, as they lack the respective C-terminal part of the G protein (470-491 aa; **Figure 5A** and Supplementary Table S5).

**Figure 5:**
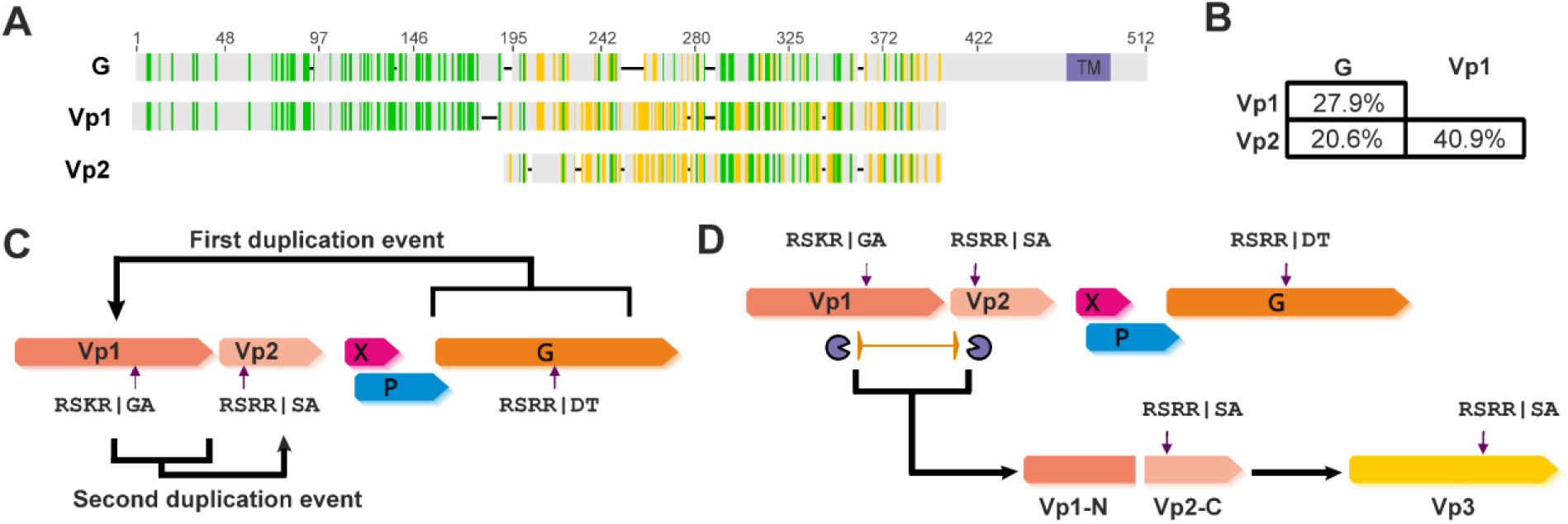
Little skate bornavirus encodes two proteins that may be the result of an ancient duplication event of the glycoprotein. **(A)** The amino acid alignment of LSBV viral proteins 1 and 2 (Vp1 and Vp2) together with the glycoprotein (G) shows, that they share homology. Vp1 and Vp2 lack the corresponding transmembrane domain (TM) of G, but each contain a predicted furin protease cleavage site (highlighted by scissors symbol). (B) Pairwise amino acid identities indicate, that Vp1 and Vp2 are more closely related to each other than to G. Therefore, in (C), supported by phylogenetic analysis (see also the Supplementary Figure S3), we hypothesised that Vp1 was first duplicated from G, followed by a second duplication of Vp1, which gave rise to Vp2. Predicted cleavage sites are indicated by arrows. (D) Transcriptional profiling suggested the possibility of alternative splicing of Vp1 and Vp2, resulting in a hybrid of the Vp1 C-terminus, including its cleavage site, and the Vp2 N-terminus, tentatively named Vp3*.

A phylogenetic tree was constructed based on an amino acid alignment of glycoproteins from selected members of the *Bornaviridae* family, supplemented by LSBV Vp1 and Vp2 (see Supplementary Figure S2). The tree provided evidence for the occurrence of a duplication event of the LSBV G gene, with Vp1 sharing the last common ancestor with G and Vp2 sharing the last common ancestor with Vp1. One possible scenario could be that initially a large part of the glycoprotein gene G was duplicated to form Vp1 and later only the part encoding the C terminus of Vp1 (representing the central part of the G) was duplicated to form Vp2 (**Figure 5C**). We also predicted potential furin endoprotease cleavage sites within Vp1, Vp2 and G, following the amino acid consensus motif `‘RS(K/R)R’` (**Figure 5C** and Supplementary Table S5).

As noted above, the predicted mRNA encoding both Vp1 and Vp2, may also undergo splicing, resulting in a hybrid ORF, tentatively named Vp3* (**Figure 5D**). The potential Vp3* protein would consist of the N-terminal portion of Vp1 and the C-terminal portion of Vp2, including the protease cleavage site. Similar to Vp1 and Vp2, the potential Vp3* would lack a transmembrane domain (Supplementary Table S5).

## DISCUSSION

Knowledge on fish bornaviruses has been limited to a single full-length genome of WhSBV [8] and a partial genome of MDCBV [7]. To identify additional and more diverse fish bornaviruses, we used an *in silico* data mining approach that screened publicly available SRA raw sequence datasets from fish (Osteichthyes and Chondrichthyes) samples. Using a similar approach, we had previously successfully identified and characterised two novel snake orthobornaviruses, CWBV and MRBV, as well as novel EBLs in reptile datasets [15]. Here, the screening combined with *de novo* assembly led to the identification of five putative complete bornavirus genomes from different samples.

We found additional sequences of WhSBV (87.9% nt identity to the previously published sequence) and MDCBV (99.5% nt identity) in fish other than the originally reported host species. The first full-length genome sequence of MDCBV presented here matched that of WhSBV in overall structure, sequence identity, and length, indicating that MDCBV and WhSBV are closely related. According to the criteria defined by the ICTV *Bornaviridae* Study Group [5], they are thought to be viruses of the same virus species (*Cultervirus hemicultri*). WhSBV was previously identified by RNA sequencing of the gut, liver, and gill tissues from a sharpbelly or wild carp (*Hemiculter leucisculus*; family Cyprinidae) from China [8], whereas MDCBV was discovered in a liver and gill tissue pool of a common carp (*Cyprinus carpio*; family Cyprinidae) during a meta-transcriptomic survey of freshwater species in the Murray-Darling Basin in Australia [7]. Here, we identified WhSBV in multiple datasets from cell lines derived from the kidney and liver of a grass carp (*Ctenopharyngodon idella*; family Cyprinidae) and MDCBV in a dataset from goldfish (*Carassius auratus*; family Cyprinidae) brain samples. Both, WhSBV and MDCBV thus appear to be members of a group of bornaviruses that are particularly common in fishes of the family Cyprinidae. Cyprinidae includes a wide range of carp and is an ancient evolutionary lineage [27]. With a global production of ∼30 million tonnes [28], carps are of great economic interest and are often cultivated in large-scale aquaculture farms. Therefore, the impact of these bornaviruses on animal health needs to be carefully assessed and the genome sequences identified in this study may provide valuable information to further investigate the distribution and variability of these viruses.

Using the data mining approach, identical WhSBV genomes were identified in datasets from the grass carp cell lines CIK (kidney) and L8824 (liver). Both cell lines originate from the Freshwater Fisheries Research Center of Chinese Academy of Fishery Sciences (formerly the Yangtze River Fisheries Research Institute) [29]. The CIK and L8824 cell lines have been repeatedly used to study viral transcriptional changes during infection, e.g. with grass carp reovirus (GCRV), and immune regulation. The presence of WhSBV in samples labelled ‘mock infection’ or ‘cell control’ (see Supplementary Table S4) indicates that both cell lines may be persistently infected and allexperimental results from experiments should be interpreted with caution. It remains unclear whether the WhSBV found in these cell lines originated from the individual(s) from which the two cell lines were derived, or whether both cell lines may have been subsequently contaminated.

We also identified four additional bornaviral genomes in non-cyprinid ray-finned fishes, and one in a cartilaginous fish. These viruses were related to WhSBV and MDCBV, but formed clearly separate taxonomic entities based on a phylogenetic analysis of N, G and L protein sequences. Despite clear differences at the nucleotide and amino acid level, four of these viruses shared the same overall genomic structure with the culterviruses WhSBV and MDCBV, and with the viruses of the genus *Carbovirus* [6]. The genome arrangement of reptilian carboviruses and these novel fish bornaviruses is peculiar in that it does not follow the standard N-X-P-<u>M-G</u>-L pattern of mononegaviruses in general and of orthobornaviruses in particular. This could indicate that the N-X-P-<u>G-M</u>-L genome arrangement evolved independently in reptile and fish bornaviruses, or that they share an ancient common ancestor that already had this genome architecture. However, assuming a virus/host co-evolution, the question arises as to why orthobornaviruses (hosts: birds, mammals, reptiles) have conserved the typical N-X-P-M-G-L pattern of mononegaviruses, although they should be comparatively younger than culterviruses (hosts: fish). It is therefore reasonable to assume that bornavirus evolution did not follow a strict virus/host co-evolution and that ancient ancestors of orthobornaviruses infected a wider range of vertebrates than extant orthobornaviruses, as it has been suggested by analysis of endogenous bornavirus-like elements (EBLs) [30]. Orthobornaviruses may therefore represent a more ancient lineage of bornaviruses, as they show the typical N-X-P-M-G-L pattern of mononegaviruses, while the N-X-P-G-M-L pattern of carbo- and culterviruses may be a more recent development.

The rearrangement of G and M may have resulted in a favourable regulation of gene expression for these viruses. By analysing the transcriptional profiles of the novel fish bornaviruses, we found that X/P and G are most likely co-expressed from the same polycistronic mRNA and M is transcribed from a spliced mRNA. In contrast, orthobornaviruses express X and P from a bicistronic mRNA starting from transcription start site S2, whereas M and G are expressed from different splice variants of mRNAs starting from S3 [31].

Genomic rearrangements do not seem to be an isolated event in bornaviruses, as illustrated by the unique genome architecture of LSBV, which encoded two more possible ORFs (Vp1 and Vp2). Both appeared to be the result of at least two independent duplication events: First, a large part of the G gene appears to have been copied into the intergenic region between N and X/P, forming Vp1. Subsequently, a part of Vp1 was duplicated into the intergenic region between Vp1 and X/P, forming Vp2 (**Figure 5**). Comparable duplication events in RNA viruses are considered very rare [32], but have been have been reported for other mononegaviruses, such as rhabdoviruses [32– 35]. Exceptionally long branches in the phylogenetic analysis indicated an accelerated evolution for Vp1 and Vp2 after the duplication events, possibly as a result of changing evolutionary context and selection pressure [36].

In addition, the Vp1 and Vp2 genes may produce a hybrid gene product Vp3* by alternative splicing, further extending the coding potential of LSBV even further. The function of Vp1, Vp2 and the splice hybrid Vp3* is currently unknown, but conserved furin cleavage sites suggest that these proteins undergo some form of post-translational modification, similar to the glycoprotein of other bornaviruses [37]. As Vp1, Vp2 and Vp3* lack a detectable transmembrane domain, it can be speculated that they may could function as soluble glycoproteins, similar to that of vesicular stomatitis virus [38]. The predicted cleavage site within the Vp1, Vp2, and Vp3* sequences may have functional significance for the virus, and future experimental investigations are needed to gain deeper insights into the unique genome architecture of this bornavirus. It would be very interesting to investigate whether other bornaviruses from cartilaginous fish share this unique genome structure, or whether LSBV is the result of an isolated evolutionary event.

Based on the phylogenetic analysis and PASC, we propose that LSBV does not belong to any of the existing genera within the family *Bornaviridae*. We have therefore submitted a taxonomic proposal to the ICTV to establish a new genus within this family. In this proposal, EEBV and FPBV were also tentatively classified as four new species within the genus *Cultervirus*.

Although the combination of gene arrangement, expression profile and potential hosts was plausible for these potential viruses, it cannot be excluded that these genomes were based on contaminated samples or inaccurate datasets and therefore did not originate from the reported host species. However, the identification of known viruses such as WhSBV and MDCBV in fish datasets related to the originally reported host, may support the credibility of our findings. Confirmation by standard methods, such as PCR and virus isolation, using independent samples from the same species would nevertheless be required to fully confirm the existence of these interesting new viruses in the reported hosts.

## CONCLUSION

Until now, WhSBV and the closely related MDCBV were the only viruses in the genus *Cultervirus*. Screening of 116,078 fish datasets from the SRA led to the identification of six tentative cultervirus genomes, including a variant of WhSBV and the first complete genome of MDCBV. These viruses may primarily infect carp, as all variants of WhSBV and MDCBV have so far been found in cyprinid samples. In addition, BDBV, EEBV, PMBV, and FPBV were discovered in a dataset from a Bombay duck, an electric eel, a Pará molly and a finned pufferfish, respectively. They had comparatively longer genomes, they all shared the same genome organisation 5’-N-X/P-G-M-L-3’, similar transcriptional profiles and regulatory sites, thus, suggesting a common ancestor of these fish bornaviruses. In addition, LSBV was identified in a little skate dataset and showed a distinct genome organisation with two additional genes that may be the result of ancient duplication events of the glycoprotein gene. The LSBV had the largest genome length (11,090 nt) of any bornavirus known to date and, to our knowledge, the presence of duplicated genes within a virus of the *Mononegaviridae* family is quite unique and has so far only been reported for a few rhabdoviruses.

The study demonstrates the power of *in silico* SRA data screening and its ability to advance the knowledge of viral diversity and evolution. The screening can easily be applied to the discovery of novel viruses from other viral families, or to the identification of known viruses in datasets from previously unknown potential host species.

## ACKNOWLEDGEMENT

We are grateful to the global scientific community for their invaluable contribution in sharing raw sequencing data. These datasets, originally intended for specific research purposes, contain valuable information that goes beyond their original purposes. We would also like to thank Robin Garcia, Jörg Linde and Michael Weber for their help with the ‘SRAminer’ pipeline.

## DATA AVAILABILITY

Nucleotide sequence data reported are available in the Third Party Annotation Section of the DDBJ/ENA/GenBank databases under the TPA accession numbers: BK063517-BK063521, BK063657 and BK063658.

