## Supplementary Figures for "Diving Deep into Fish Bornaviruses: Uncovering Hidden Diversity and Transcriptional Strategies through Comprehensive Data Mining"

| | LSBV BK063518 | PMBV BK063657 | BDBV BK063658 | FPBV BK063517 | EEBV BK063519 | WhSBV NC_055169 | WhSBV BK063520 | MDCBV BK063521 |
|---|---|---|---|---|---|---|---|---|
| **little skate bornavirus BK063518 (LSBV)** | 100.0 | 38.4 | 39.9 | 39.2 | 39.6 | 38.4 | 39.2 | 38.2 |
| **Pará molly bornavirus BK063657 (PMBV)** | 38.4 | 100.0 | 65.8 | 44.2 | 43.0 | 43.7 | 42.1 | 42.9 |
| **Bombay duck fish bornavirus BK063658 (BDBV)** | 39.9 | 65.8 | 100.0 | 44.7 | 42.5 | 43.9 | 44.3 | 43.5 |
| **finepatterned puffer bornavirus BK063517 (FPBV)** | 39.2 | 44.2 | 44.7 | 100.0 | 44.6 | 44.1 | 44.3 | 43.7 |
| **electric eel bornavirus BK063519 (EEBV)** | 39.6 | 43.0 | 42.5 | 44.6 | 100.0 | 49.7 | 50.3 | 51.5 |
| **Wǔhàn sharpbelly bornavirus NC_055169 (WhSBV)** | 38.4 | 43.7 | 43.9 | 44.1 | 49.7 | 100.0 | 87.9 | 78.1 |
| **Wǔhàn sharpbelly bornavirus BK063520 (WhSBV)** | 39.2 | 42.1 | 44.3 | 44.3 | 50.3 | 87.9 | 100.0 | 78.5 |
| **Murray-Darling carp bornavirus BK063521 (MDCBV)** | 38.2 | 42.9 | 43.5 | 43.7 | 51.5 | 78.1 | 78.5 | 100.0 |

**Supplementary Figure S1:** Selected PASC nucleotide identities (%) based on complete genomes.

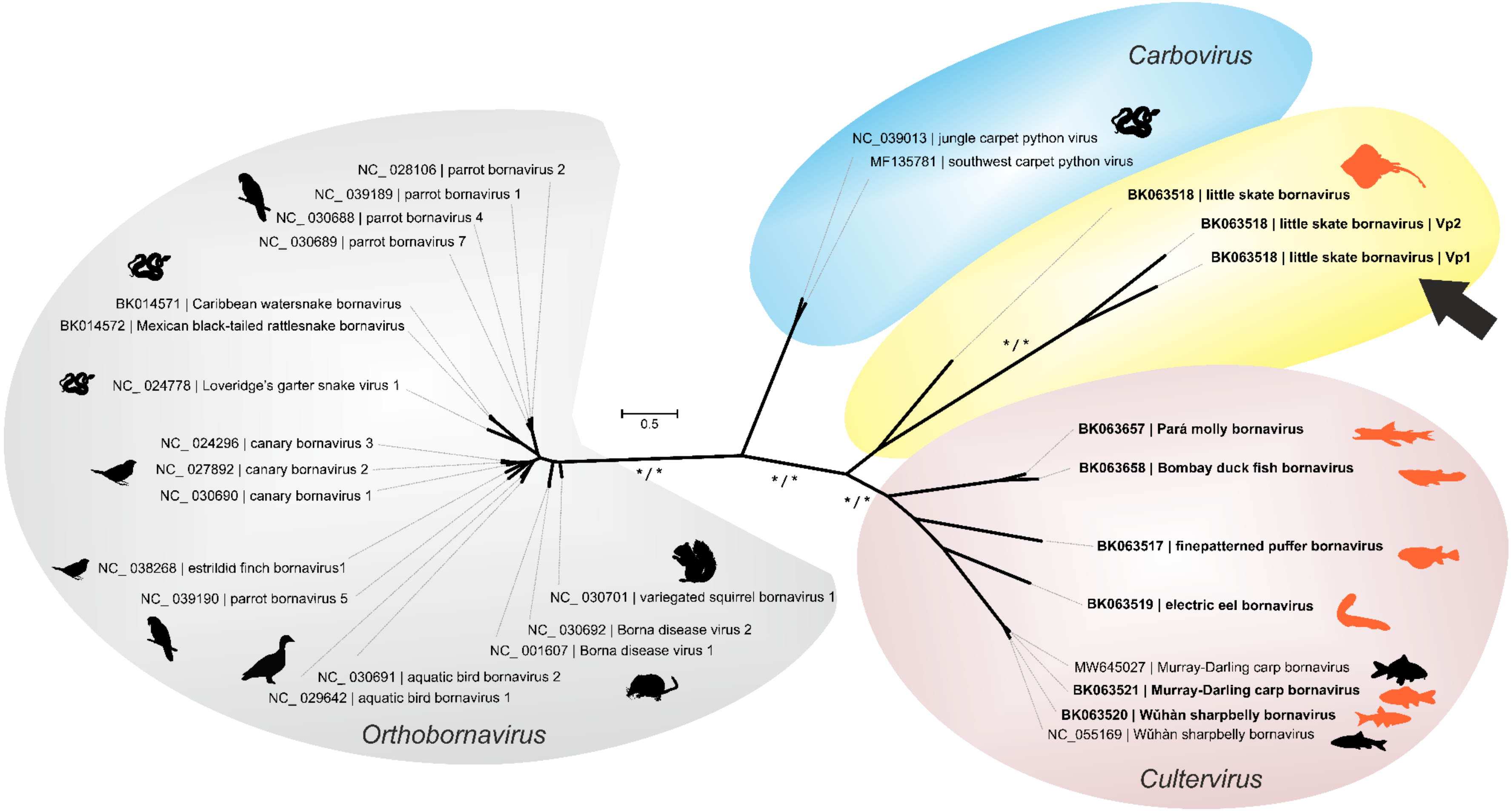


**Supplementary Figure S2: Phylogenetic relationships within the family *Bornaviridae* based on the G protein.** The maximum-likelihood tree was based on the amino acid sequence alignment (total length of 596 aa) of the viral protein G of the potential fish bornaviruses (bold) together with representative members of the genera *Cultervirus*, *Carbovirus* and *Orthobornavirus*. The glycoprotein of little skate bornavirus and the predicted Vp1 and Vp2 (arrow) share a last common ancestor, indicating ancient duplication events. Note the comparably long tree branches towards Vp1 and Vp2. The silhouettes represent typical host organisms of previously published bornaviruses or the reported sampling source (highlighted) of the viral genomes identified in this study. Sequences from this study are shown in bold. The tree was constructed using IQ-TREE (version 2.2.2.3), an optimal partitioning model and statistical support of 1 million replicates each for the ultrafast bootstrap and SH-aLRT tests. Statistical support is shown for major branches.

**Supplementary Table S1:** See file ”Supplementary Table S1.xlsx”

**Supplementary Table S2:** See file ”Supplementary Table S2.xlsx”

**Supplementary Table S3:** See file ”Supplementary Table S3.xlsx”

**Supplementary Table S4:** See file ”Supplementary Table S4.xlsx”

**Supplementary Table S5:** See file ”Supplementary Table S5.xlsx”
